# De novo Engineered Living Materials via Elastin-Like Polypeptide-Mediated Self-Assembly

**DOI:** 10.1101/2025.06.26.661578

**Authors:** Sarah B. Browning, Davide Prati, Luca Mascia, Paolo Magni, Lorenzo Pasotti, Sara Molinari

## Abstract

De novo engineered living materials (ELMs) are cellular systems that self-assemble into macroscopic structures through genetically encoded interactions, offering a route to programmable materials grown directly from living cells. Despite their promise, the molecular design principles that enable scalable self-assembly in de novo ELMs remain poorly understood. Here, we engineer *Escherichia coli* to display elastin-like polypeptides (ELPs) on the outer membrane, transforming single cells into self-assembling living materials through weak intermolecular interactions. By tuning the polarity of ELP sequences, we generate assemblies spanning micrometer- to centimeter-length scales that sediment within a few hours while preserving cellular metabolic activity. We demonstrate the portability of this platform across genetic backgrounds and inducible expression systems, and deploy it in an ethanologenic *E. coli* strain as a proof of principle. In small-scale fermentation settings, ELP-based ELMs enable controllable flocculation, reduce filtration time by more than threefold, and maintain ethanol production performance comparable to that of the parental strain. Together, this work establishes ELP surface display as a modular strategy for constructing de novo engineered living materials and defines initial genetic design rules linking molecular-scale interactions to emergent macroscopic organization.

## Introduction

Engineered living materials (ELMs) are composites of cells embedded in a biopolymer matrix, which assembles them into macroscopic multicellular structures^1^. Because they incorporate living cells, they hold the promise to revolutionize the field of material science by enabling metabolically active and environmentally-responsive matter^2^ whose exact behaviors can be rationally programmed using synthetic biology approaches. In this way, living materials could be designed to continuously produce certain molecules^3^ or specifically sense and respond to tailored inputs^4,5^. By supporting the self-assembly of macroscopic cellular structures, ELMs could provide the advantages of natural biofilms^6,7^, such as improving biological functions by protecting inner bacteria and favoring metabolic cooperations^8^. Moreover, by virtue of being macroscopic, they can be more easily handled than planktonic cells, potentially allowing for bacterial deployment in real-world, field-relevant applications^5,9,10^. In addition, ELMs could spatially confine and concentrate cells for improved biocontainment and spatial separation^11^. Since living cells serve as renewable, biodegradable building blocks, they also have the potential to improve sustainability and manufacturing costs compared to standard materials^1^.

So far, ELMs have been manufactured using different strategies, involving mixing external matrices with bacterial cells^12,13,5,14,15^, or processing microscopic cellular assemblies into larger-scale materials^16–18^. Alternatively, de novo ELMs are obtained from bacterial cells genetically engineered to produce the biopolymer matrix responsible for their self-assembly^19–21^. In this way, de novo ELMs grow from progenitor cells, without the need for specialized equipment or external processing and with minimal energy input, resembling the process of generation of natural living materials, such as wood, skins, or biofilms^1^. Due to cells’ ability to produce the materials, de novo ELMs have the potential to regenerate and self-repair in case of damage, maximizing robustness and longevity and minimizing maintenance costs.

To obtain de novo ELMs, bacteria must be engineered to produce an extracellular matrix able to assemble cells into cohesive multicellular structures. Cellular self-assembly has been obtained by genetically programming microorganisms to display adhesins forming strong pair-wise interactions, such as coiled coils^20^, nanobody-antigen pairs^21^, and SpyTag-SpyCatcher^8^. These designs have shown the formation of bacterial assemblies whose size could be tuned by modulating the strength of adhesin interactions^20^. However, de novo ELMs produced in this way never exceeded the microscopic scale, with cell clusters reaching at most hundreds of micrometers^20,21^. More recently, work in *Caulobacter crescentus* has shown that displaying peptides that form multiple weaker hydrophobic interactions can drive larger-scale assembly, up to centimeters^19,22^. This observation is consistent with findings on DNA-coated nanoparticles^23^, where interparticle links that can readily break and reform enable the system to explore lower-energy conformations, ultimately supporting the formation of larger crystals. Despite providing an important proof of principle, this study has several limitations as a general platform for elucidating the design rules governing weak interaction-driven bacterial self-assembly, primarily due to the limited genetic tractability of *C. crescentus* and its distinctive surface architecture^19^.

To overcome this challenge, we engineered the model bacterium *Escherichia coli* to display elastin-like polypeptides (ELPs). ELPs are synthetic peptides derived from the repetitive sequence of the human tropoelastin^24^ known to form multiple weak hydrophobic interactions among their chains. They consist of pentameric repeats of Valine-Proline-Glycine-X-Glycine where X represents a guest residue that can be any amino acid except proline^22^. At a constant temperature, the ELP interactions are primarily governed by the amino acid sequence, i.e., the identity of the guest residue (X) and the number of repeats^22,25,26^. Thus, the tunable sequence of ELPs and their well-established aggregation make them an exceptionally tractable experimental system for probing how molecular-level interactions of surface-displayed peptides direct bacterial self-assembly.

The model bacterium *E. coli* represents an ideal chassis because, unlike *C. crescentus*, it supports peptide surface display without the need for large secretion tags that could bias intermolecular interactions^19^. Furthermore, *E. coli* possesses extensive genetic tools enabling both stable and inducible protein expression from plasmid DNA, which simplifies the screening of multiple peptide sequences. By tuning the ELP sequence and mapping the resulting changes in bacterial self-assembly, this platform enabled us to define initial engineering principles for the design of de novo flocculating *E. coli*-based engineered living materials (FloEc-ELMs).

As proof of principle, we demonstrated the utility of FloEc-ELM formation for the downstream processing of microbial fermentations. Downstream processing, particularly biomass separation, remains a major bottleneck in microbial bioproduction due to its high energy demand, capital cost, and susceptibility to membrane fouling^27–32^. Conventional approaches such as centrifugation and cross-flow filtration are infrastructure-intensive, limiting their scalability and reuse in continuous or reinoculation-based processes^29^. Microbial self-assembly—also known in the fermentation context as flocculation—offers an attractive strategy to simplify biomass separation, promoting rapid cell sedimentation, thereby reducing reliance on energy-intensive mechanical or membrane-based methods^32,33^. While naturally flocculating yeasts have been successfully exploited in industrial fermentations, common production hosts, such as *E. coli,* lack this capability^34^. Existing genetic, chemical, and physical approaches to induce flocculation in *E. coli* improve broth clarification but introduce significant drawbacks, including metabolic burden, limited portability, added costs, and process contamination^35,36^. As a result, there remains a need for a broadly applicable, low-burden strategy to enable controllable biomass aggregation in microbial bioproduction.

Here, we present tunable self-assembly of FloEc-ELMs as a generalizable strategy for inducing controllable flocculation and demonstrate its utility as a proof of principle for improving biomass separation in a microbial fermentation process focused on lactose wastewater valorization. Specifically, we transferred the FloEc-ELM technology into an ethanol-producing strain of *E. coli* and observed that this genetic modification can improve bacterial biomass recovery during the cell separation step at the end of fermentation, with a significant reduction of filtration time for the fermented broth after sedimentation, and without compromising the ethanol production performance. Sedimented cells were also viable and could be successfully used to start a new lactose-to-ethanol fermentation in a dairy effluent.

## Results

### 2.1 Formation of FloEc-ELMs via Surface Display of ELPs in E. coli

We engineered FloEc-ELMs by displaying elastin-like polypeptides (ELPs) on the surface of *E. coli*. Previous studies have shown that partially substituting the canonical valine guest residues in ELP sequences with alanine enhances protein expression^37^ and surface display in *E. coli*^18^. In addition, alanine-rich ELPs were reported to form cohesive bacterial films upon cell concentration by vacuum filtration, suggesting a potential role in mediating cell-cell adhesion^18^. However, because the formation of multicellular structures in these studies required external processing that forced cell compaction, it remained unknown whether ELP surface display alone is sufficient to drive spontaneous, self-directed bacterial assembly in suspension. To verify the ability of alanine-rich ELPs to drive microbial self-assembly, we cloned a 10-repeat ELP construct, containing six alanine substitutions (referred to as ELP_10_-ALA), fused to the EhaA autotransporter for extracellular display (Fig. 1A)^20^. For characterization purposes, we included a SpyTag^38^ and a HisTag^39^ to the opposite termini of the ELP sequence. This construct was placed under the control of an IPTG-inducible promoter (T5/lac), which allows for the fine regulation of gene expression in *E. coli*^40^.

**Figure 1.**
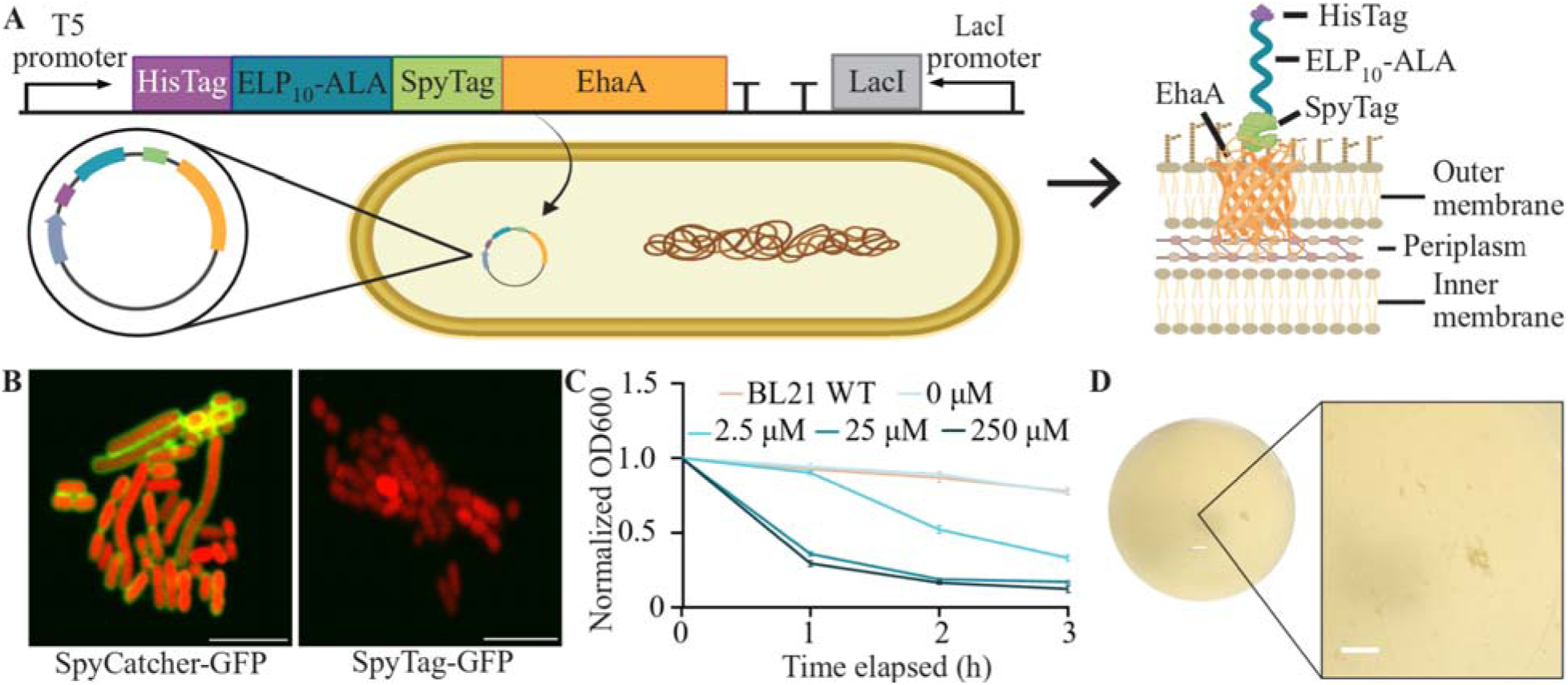
Surface display of ELP_10_-ALA yields FloEc-ELMs. A) Schematic of the ELP_10_-ALA expression cassette including the T5/lac inducible promoter, the EhaA autotransporter, and the HisTag and SpyTag, respectively at the N and C terminus of the ELP_10_-ALA sequence^20^. B) Confocal microscopy images of *E. coli* cells displaying ELP_10_-ALA stained with SpyCatcher-GFP (left panel) or SpyTag-GFP (right panel). Scale bars are 5 µm. C) Sedimentation assay of ELP_10_-ALA FloEc-ELMs induced with increasing concentrations of IPTG. Error bars are plotted around the mean of each time point and represent 95% confidence intervals of six samples. D) Representative photograph of a flask bottom containing FloEc-ELMs originated from the self-assembly of *E. coli* cells displaying ELP_10_-ALA. Additional images can be found in Supplementary Fig. 4. Scale bars are 0.5 cm.

The expected extracellular localization of ELP_10_-ALA was confirmed via confocal microscopy (Fig. 1B) after staining the cells with SpyCatcher-GFP. SpyCatcher-GFP specifically bound the SpyTag included in the displayed construct^19^, resulting in fluorescent halos around bacteria expressing intracellular mScarlet (red fluorescent protein) from a separate plasmid (Fig. 1B—left panel). In contrast, cells incubated with SpyTag-GFP (Fig. 1B—right panel)—as a negative control—do not show any green fluorescent staining, confirming that the extracellular fluorescence observed is due to specific SpyCatcher-SpyTag binding. Western blot analyses (Supplementary Fig. 1), performed with anti-HisTag antibody, validated the integrity and correct molecular weight of ELP_10_-ALA (∼46.17 kDa), with strong enrichment in the outer membrane fractions, consistent with surface display. Flow cytometry (Supplementary Fig. 2) provided quantitative confirmation of these findings, showing significantly—two orders of magnitude—higher fluorescence intensity in SpyCatcher-GFP-stained samples compared to SpyTag-GFP.

To evaluate the effect of ELP_10_-ALA display on cell viability, we incubated cells with propidium iodide (PI), a dye that selectively stains cells with compromised outer membranes^41^. Confocal microscopy (Supplementary Fig. 3) revealed that ELP_10_-ALA surface display substantially reduced viability, evident from a significantly higher proportion of red-fluorescent cells compared to the wild-type strain. This observation was further supported by the frequent appearance of elongated cell morphologies, indicative of cellular stress.

To evaluate the ability of displayed ELP_10_-ALA to mediate the formation of FloEc-ELMs—resulting from cellular self-assembly—we conducted sedimentation assays (Fig. 1C) to quantify deposition kinetics. Cells were cultivated in shaking conditions and induced during mid-log phase (OD_600_ 0.4–0.6). Two hours after the induction, we incubated the cells statically to allow for deposition. To determine the optimal expression level of ELP_10_-ALA, we compared sedimentation at various IPTG concentrations. Results (Fig. 1C) show that the most efficient sedimentation occurred at 25 µM IPTG, while increasing the inducer concentration to 250 µM provided no significant further benefit. For this level of induction, complete settling of ELMs was achieved after 2 hours of static incubation, with a final OD of 0.188, in contrast to the higher residual ODs of 0.892 and 0.870 observed for non-induced and non-transformed controls, respectively. To visually assess the dimensions of FloEc-ELMs resulting from cellular self-assembly, we imaged the bottom of the flask where they formed after an overnight growth. Results (Fig. 1D and Supplementary Fig. 4) show that, despite the rapid cell depositions, the cluster size of FloEc-ELMs remains microscopic, with only sporadic millimeter-sized structures occasionally appearing in the liquid cultures.

Taken together, these results confirm that ELP_10_-ALA surface display drives the formation of FloEc-ELMs that enhance sedimentation efficiency relative to wild-type cells and appear a sporadic macroscopic structures in liquid cultures. This genetic modification is accompanied by a notable reduction in cell viability, presenting a critical trade-off for ELM applicability.

### 2.2 Polarity-Driven Optimization of ELP Sequences Enhances Viability and Enables Scalable FloEc-ELM Formation

To promote the formation of macroscopic FloEc-ELMs beyond the sporadic assemblies observed in cells displaying ELP_10_-ALA, we tuned ELP sequence polarity. This design choice was motivated by evidence that individual ELP chains immobilized on surfaces can adopt collapsed conformations^42^. We hypothesized that hydrophobic ELPs are more prone to chain collapse, which could hinder displayed chain interlocking, reducing their ability to mediate cell-cell interactions and thereby limit FloEc-ELM formation. To test this hypothesis, we designed two alternative ELP variants in which alanine was substituted with polar residues: either positively charged lysine (ELP_10_-LYS) or negatively charged glutamic acid (ELP_10_-GLU).

We first evaluated surface display by staining the cells with SpyCatcher-GFP, followed by flow cytometry quantification. Negative control-normalized GFP intensities (Fig. 2A and Supplementary Fig. 5) indicate that, although all constructs exhibit display levels within the same order of magnitude, ELP_10_-GLU shows more than a twofold increase in surface display relative to ELP_10_-ALA, whereas ELP_10_-LYS displays 42.8% lower fluorescence than the alanine-substituted construct.

**Figure 2.**
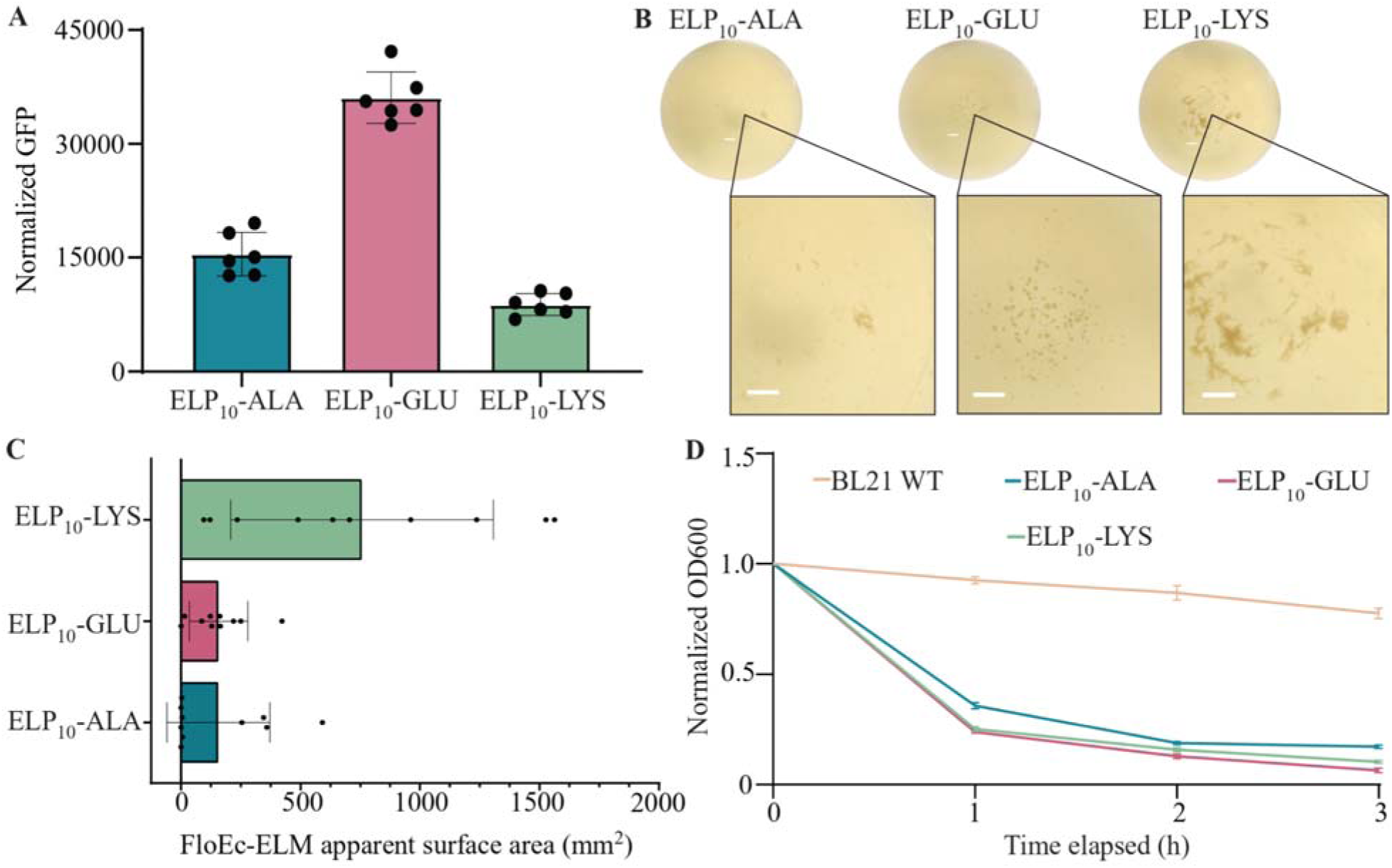
ELP polarity affects cell viability and FloEc-ELM production. A) Flow cytometry of the ELP-displaying cells stained with SpyCatcher-GFP and normalized on SpyTag-GFP. Error bars are centered on the mean value and represent 95% confidence intervals of six samples. B) Photographs of FloEc-ELMs grown in Erlenmeyer flasks, showing macroscopic FloEc-ELMs Scale bars are 0.5 cm. The panel relative to ELP_10_-ALA is the same as in Fig. 1D, included here to facilitate direct comparison. C) Image quantification of macroscopic FloEc-ELMs apparent surface area, defined as the pixels contained in the assembly mask multiplied by the squared conversion factor of pixels to millimeters. Error bars are centered on the mean value and represent 95% confidence intervals of ten samples. D) Sedimentation assays comparing the deposition rates of different FloEc-ELMs induced with 25 μM IPTG. ELP_10_-LYS showed a significant reduction in OD_600_ compared to ELP_10_-ALA (p=0.0001) and ELP_10_-GLU (p=0.0043) after the first hour. Error bars are centered on the mean value and represent 95% confidence intervals of six samples.

Remarkably—in contrast to ELP_10_-ALA—bacteria displaying either ELP_10_-LYS or ELP_10_-GLU formed macroscopic FloEc-ELMs more consistently across different cultures, extending up to several millimeters in length (Fig. 2B—central and right panels, and Supplementary Fig. 6). Among the polar variants, ELP_10_-LYS yielded the largest observed FloEc-ELMs, with the biggest living material pieces reaching approximately 1.75 cm in length (Supplementary Fig. 7). Bacteria displaying ELP_10_-GLU equally formed millimeter-sized particles (Supplementary Fig. 6), albeit at a much lower concentration compared to ELP_10_-LYS and with a slightly different circular morphology.

To quantitatively compare the total amount of macroscopic FloEc-ELMs that each sequence is able to produce, we calculated the total FloEc-ELMs apparent surface area^19^ (Fig. 2C) through image analysis of the flask bottom photographs by quantifying the pixels occupied by FloEc-ELMs. While this approach does not account for the three-dimensional thickness of the living materials, it allows for the quantification and comparison of different FloEc-ELMs directly in the flasks where they form. This method is therefore advantageous because it avoids the sampling bias and sample disruption imposed by microscopy and the indirect measure of the sedimentation assay.

Sedimentation assays reflect the trend emerged in the image analysis, showing that both polar constructs facilitated more efficient cell deposition than ELP_10_-ALA, as indicated by a final OD of 0.17 and 0.158 for ELP_10_-GLU and ELP_10_-LYS respectively—representing a 9.6% and 16% improvement over the alanine-substituted construct (Fig. 2D). However, the sedimentation assays fail to highlight the magnitude of difference in the macroscopic FloEc-ELMs production among the three displayed ELP sequences, clearly observable in the flask bottom pictures and properly captured by the FloEc-ELMs apparent surface area quantification.

PI staining—indicative of cell membrane integrity—shows substantially reduced toxicity in both polar constructs compared to ELP_10_-ALA (Supplementary Fig. 8), suggesting that the hydrophobicity of displayed peptides also contributes to cytotoxicity. Among the tested variants, ELP_10_-LYS best preserved cell viability.

Based on superior production of FloEc-ELMs and cell viability, we selected ELP_10_-LYS for further experimentation. Western blot analysis (Supplementary Fig. 1) of bacteria displaying the ELP_10_-LYS construct confirmed the expected molecular weight (∼46.52kDa) and enrichment of the outer membrane fraction, consistent with surface display.

Together, these findings underscore the advantages of polar substitutions in ELP design for enabling the formation of robust, large-scale FloEc-ELMs, while also preserving cell viability.

### 2.3 Transferring FloEc-ELM Technology to an Ethanol-Producing E. coli Strain for Tunable Flocculation in Fermentation Settings

To demonstrate the potential of FloEc-ELMs in a bioproduction process, we transferred the ELP_10_-LYS display system into the industrially relevant ethanologenic *E. coli* strain W105Fe^43^ engineered to maximize ethanol production from lactose as a carbon source. Since the original IPTG-inducible promoter would be activated by the presence of lactose in the media, we refactored the system to be controlled by an HSL/LuxR inducible promoter^44,45^. In this design, a constitutive promoter drives expression of the LuxR transcriptional regulator, which binds its cognate ligand N-3-oxohexanoyl-L-homoserine lactone (HSL) upon diffusion into the cell. The resulting LuxR–HSL complex activates transcription from the Plux promoter, thereby enabling controlled induction of ELP_10_-LYS. This system allowed us to supplement the LB media with 2 g/L lactose to simulate fermentation-relevant conditions.

To identify the optimal induction level of the ELP_10_-LYS display in the W105Fe strain, we performed sedimentation assays across a range of HSL concentrations (Fig. 3A). Results showed that the induction with 2 nM HSL efficiently triggered FloEc-ELMs formation comparably to what observed in BL21 cells induced with 25 µM IPTG (Fig. 2C). Whereas, increasing the HSL concentration to 20 or 200 nM did not further enhance sedimentation. The PI staining of the 2 nM HSL-induced W105Fe–ELP_10_-LYS (Fig. 3B—middle panel) strain excluded significant loss of viability as result of the genetic modification, indicated by levels of red fluorescence not much higher than the non-engineered strain (Fig. 3B—left panel). In contrast, cells induced with 20 nM HSL displayed a much greater proportion of PI-stained cells (Fig. 3B—right panel), suggesting that overexpression of the surface-displayed ELP_10_-LYS disrupts membrane integrity. Based on these results, we selected 2 nM HSL as the optimal induction level for subsequent experiments, balancing robust FloEc-ELM formation with preserved cell viability.

**Figure 3.**
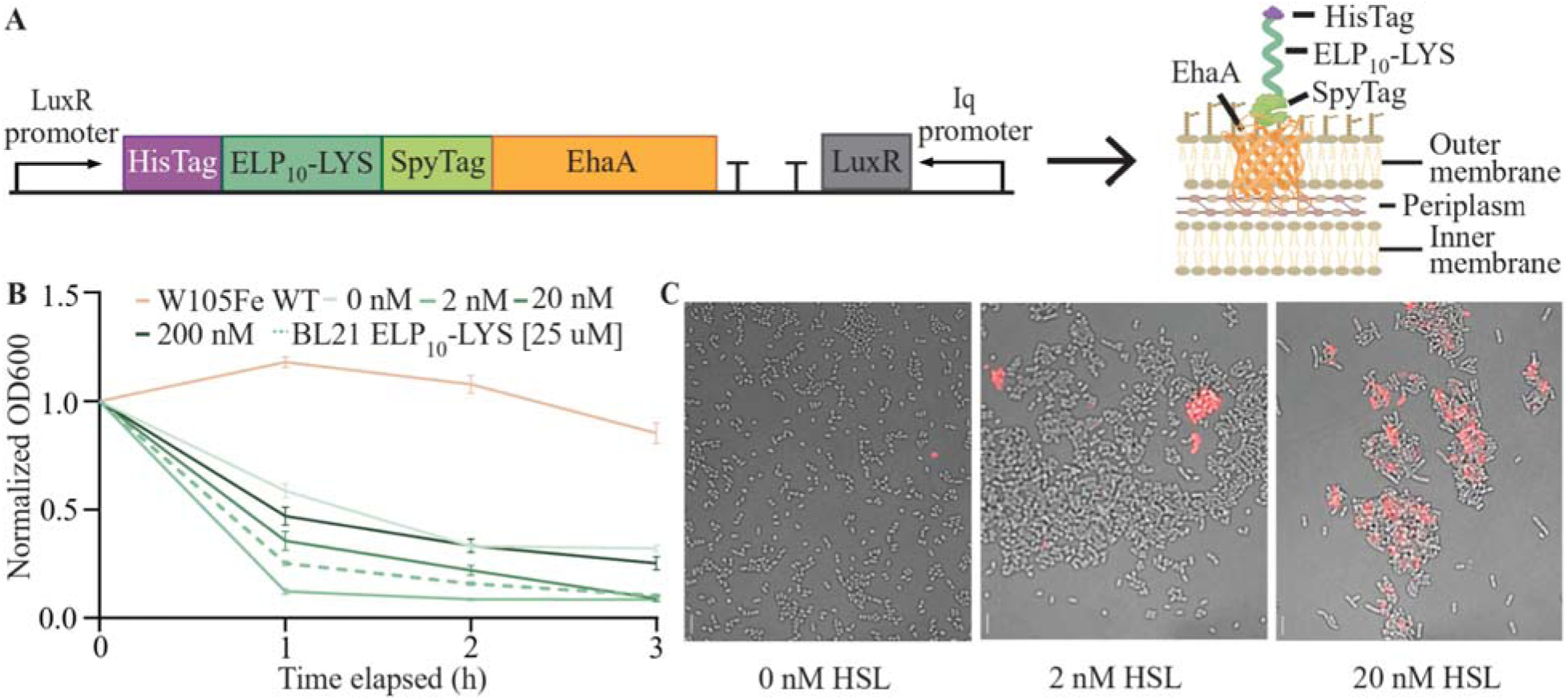
Characterization of FloEc-ELMs engineered in the ethanologenic strain W105Fe. A) Schematics of the refactored expression cassette for ELP_10_-LYS controlled by the HSL/LuxR system cloned in the W105Fe strain. B) Sedimentation assay showing the rates of FloEc-ELM deposition across the HSL induction range (0, 2, 20, and 200 nM) for the base W105Fe and the W105Fe-ELP_10_-LYS strain. The dashed line represents the ELP_10_-LYS system controlled by the T5/lac promoter and induced by 25 µM IPTG in BL21 (Fig. 2D—green line) for comparison. C) Confocal images of PI-stained cells of W105Fe-ELP_10_-LYS cells at 0 nM (left panel), 2 nM (middle panel), and 20 nM induction (right panel) showing increasing loss of cell viability. Scale bars are 5 μm.

To ensure proper extracellular display of ELP_10_-LYS in W105Fe, we stained induced W105Fe cells with SpyCatcher-GFP and measured fluorescence intensity by flow cytometry. Results (Supplementary Fig. 9) confirmed the expected extracellular localization of the construct upon induction with 2 nM HSL.

These findings demonstrate that the FloEc-ELM platform can be effectively transferred across *E. coli* strains and coupled with different inducible gene expression systems, yielding robust FloEc-ELM formation without compromising cellular viability.

### 2.4 FloEc-Elms Enhance Filtration Efficiency and Biomass Recovery in Fermentation Settings

We next evaluated the HSL-inducible ELP_10_-LYS FloEc-ELM system in the ethanologenic *E. coli* strain W105Fe during lactose fermentations, focusing on its impact on ethanol production and biomass settlement. We first evaluated whether FloEc-ELM formation could be induced at the end of the lactose fermentation, when the biomass needs to be separated from the ethanol-containing supernatant^32,46^. However, cultures induced at high cellular density (Supplementary Fig. 10)—which mimic late growth conditions—indicate limited self-assembly, likely due to nutrient depletion restricting protein expression. In contrast, rapid sedimentation occurred when cultures were induced at lower densities (Supplementary Fig. 10), suggesting that active growth and nutrient availability are prerequisites for FloEc-ELM formation. Based on these findings, in the following experiments, we induced ELP_10_-LYS expression during exponential growth at the beginning of the fermentation process.

To assess the effects of early ELP_10_-LYS induction on W105Fe’s ability to produce ethanol, we compared batch fermentation titers of the base (W105Fe) and the induced and non-induced engineered strains (W105Fe-ELP_10_-LYS). Fermentations were performed in a rich medium containing 40 g/L lactose to ensure sustained ethanol production, and cells were induced in the exponential phase.

Ethanol titers (Fig. 4A) measured at the end of fermentation (24 h) were comparable across all strains, with final concentrations of approximately 17.5 g/L and complete lactose consumption, corresponding to 80% of the maximum theoretical yield of ethanol production^47^. These results indicate that FloEc-ELM formation does not impair ethanol productivity.

**Figure 4.**
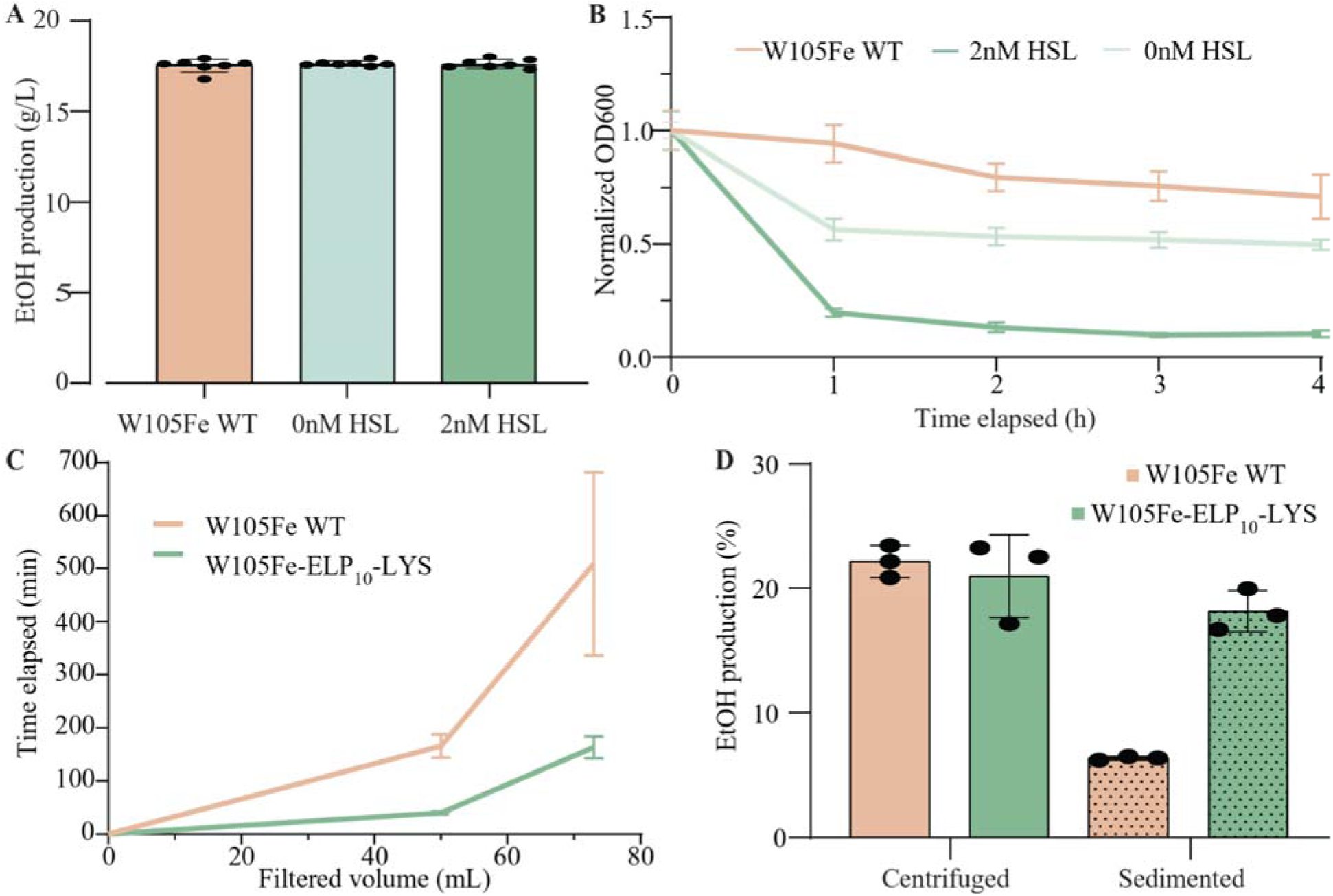
FloEc-ELMs assembly does not impair ethanol production and improves biomass recovery in a batch fermentation process. A) Ethanol production quantification for the W105Fe (parent), and induced and non-induced W105Fe-ELP_10_-LYS ethanologenic strains in LB Lennox with 40 g/L lactose, after 24 h. Error bars are centered on the mean value and represent 95% confidence intervals of seven samples. B) Sedimentation assay conducted after a 24-h batch fermentation. Error bars are centered on the mean value and represent 95% confidence interval of three samples. C) Filtered volume over time of the W105Fe (n=2) and the induced W105Fe-ELP_10_-LYS (n=3) after a 24-h batch fermentation in LB Lennox with 40 g/L lactose. D) Ethanol production quantification for the W105Fe and the induced W105Fe-ELP_10_-LYS strains in dairy waste fermentation medium after 72 h. Fermentation medium was inoculated with bacterial biomass recovered from the pre-culture by centrifugation (Centrifuged) or sedimentation (Sedimented). Error bars are centered on the mean value and represent 95% confidence intervals of three samples.

Using the same fermentation setup, we next evaluated whether rapid FloEc-ELM sedimentation could facilitate filtration of the supernatant, a required step for separating the broth from the bacterial biomass prior to ethanol recovery. Sedimentation assays showed that the induced W105Fe-ELP_10_-LYS strain achieved a 90% reduction in cell density in the supernatant after 4 h of static incubation (87% after only 2 h), whereas the non-flocculating parental strain only reached a 29% decrease (Fig. 4B). The non-induced W105Fe-ELP_10_-LYS strain exhibited an intermediate OD_600_ reduction of 50%, likely due to leaky expression from the inducible promoter. These results suggest that FloEc-ELMs—formed upon induction in the exponential phase—remain stable throughout the fermentation process, effectively enhancing biomass sedimentation.

Next, we determined the effects of improved biomass sedimentation on supernatant filtration. Filtration assays of 75 mL post-fermentation cultures allowed to sediment for 1.5 h (Fig. 4C) showed that the FloEc-ELM-forming culture has a 3.1-fold increase in filtration flow rate (26.8 mL/min) compared to the non-flocculating control (8.6 mL/min), allowing for shorter processing time. Moreover, the induced W105Fe-ELP_10_-LYS strain yielded more consistent filtration rates across replicates, as filtration failed to complete in some non-flocculating cultures due to filter clogging (N=2, data not shown). This suggests that FloEc-ELM production also enhances process robustness.

As a further demonstration of FloEc-ELM utility, we evaluated whether flocculating cells could be used directly as inoculum for fermentations of lactose-rich cheese whey permeate, a dairy industry wastewater stream commonly used as a substrate for industrial ethanol production^43^. This medium was selected because efficient lactose fermentation requires high initial cell densities^43^, which are typically achieved by concentrating precultures via centrifugation before reinoculation. To benchmark this standard approach, centrifugation-based biomass recovery was used as a control (Fig. 4D—Centrifuged), in which the entire concentrated preculture was used to inoculate whey permeate medium. In parallel, we tested sedimentation-based biomass recovery enabled by FloEc-ELM formation (Fig. 4D—Sedimented). After 1.5 h of static incubation, precultures were gently decanted until approximately 100 µL (∼10% of the original volume) remained. Under these conditions, induced W105Fe-ELP_10_-LYS cultures yielded a clearly visible sediment suitable for reinoculation, whereas the non-flocculating parental strain did not (Supplementary Fig.11).

When biomass was recovered by standard centrifugation, both W105Fe and W105Fe-ELP_10_-LYS achieved comparable ethanol titers in dairy waste fermentations (Fig. 4D—Centrifuged), reaching 22 and 21 g/L, respectively. In contrast, when biomass recovery relied on sedimentation alone, ethanol production was markedly strain dependent. Under these conditions, W105Fe–ELP_10_-LYS reached an ethanol titer of 18.1 g/L—approximately threefold higher than that of the non-flocculating parental strain (Fig. 4D—Sedimented). Based on sugar consumption, W105Fe–ELP_10_-LYS achieved similar ethanol yields under centrifugation- and sedimentation-based recovery (65% and 61% of theoretical, respectively), indicating that sedimentation primarily affects inoculum recovery rather than strain productivity.

Altogether, these findings highlight some of the practical benefits of FloEc-ELMs for industrial fermentation, including faster broth clarification, reduced mechanical stress on downstream equipment, and the potential for lower costs associated with membrane cleaning or replacement. Improved clarification also enables efficient recovery of viable biomass for reuse.

## Discussion

In this study, we show that de novo engineered living materials can be generated by genetically programming *E. coli* to display elastin-like polypeptides (ELPs) on their outer membrane, thereby converting planktonic cells into self-assembling, gravity-settling multicellular materials (FloEc-ELMs). Unlike previous adhesion-based systems that rely on strong, pairwise interactions and remain confined to the microscale, ELP-mediated interactions enable the emergence of macroscopic living materials spanning millimeters to centimeters. This work establishes ELP surface display as a minimal and genetically encoded strategy for driving large-scale bacterial self-assembly.

Tuning of ELP sequence composition revealed that peptide polarity is a key determinant of both host compatibility and emergent material properties. Substitution of hydrophobic alanine residues with charged amino acids significantly improved cell viability while simultaneously promoting the formation of larger and more robust FloEc-ELMs. Notably, the positively charged ELP_10_-LYS variant generated the largest assemblies despite exhibiting the lowest surface display levels, suggesting that maximal ELP adhesion strength or expression is not required for macroscopic material formation. Instead, weaker, reversible interactions—potentially mediated by electrostatics and/or changes in the effective presentation of ELP chains at the cell surface—may allow better interlocking of ELP chains and dynamic bond rearrangement that enables structural reorganization and growth into larger assemblies.

The formation of FloEc-ELMs across distinct *E. coli* backgrounds demonstrates that ELP-mediated self-assembly is portable and not restricted to a single host physiology. Moreover, refactoring the expression from IPTG control to a LuxR/HSL induction system shows that FloEc-ELMs can be readily rewired into different genetic expression systems, with host-specific factors primarily tuning assembly strength and morphology rather than enabling it. Importantly, ELP-based ELM formation seems to impose minimal metabolic burden, as fermentation data show that ethanol productivity remains largely unperturbed by surface display.

As proof of principle, we show that de novo ELM formation can be harnessed to simplify downstream processing in microbial fermentations. FloEc-ELMs enable rapid biomass sedimentation and yield a more than threefold increase in filtration efficiency, translating into reduced membrane fouling, improved process robustness, and potential energy savings. Beyond ethanol production, this strategy could be broadly applicable to bioprocesses where products are extracellularly secreted, where biomass reuse is desired, or where physical containment of genetically modified organisms is required before downstream purification. Moreover, the ability to recover viable biomass via sedimentation alone enables centrifugation-free reinoculation, as demonstrated here for lactose-rich dairy waste fermentations.

More broadly, this work establishes a genetically tractable platform for elucidating design principles governing macroscopic self-assembly in de novo ELMs. By varying ELP sequence polarity, length, and composition, future studies can systematically map molecular-scale interactions to emergent material properties. Coupled with functional protein domains and alternative—non model—bacterial hosts, this approach opens new avenues for constructing programmable, self-assembling living materials with tunable mechanical, physical and biological properties.

## Methods

### 4.1 Bacterial Strains

The strains used in this work for characterizing ELMs are derivatives of *E. coli* BL21—referred to as wild-type. For fermentation experiments and FloEc-ELM formation under ethanol production conditions, the W105Fe ethanologenic strain was used^43^. W105Fe is a derivative of *E. coli* W (DSM 1116), engineered for optimized ethanol production from lactose-rich dairy waste by i) the chromosomal insertion of a constitutive codon-optimized adhB-pdc operon (coding for an alcohol dehydrogenase II and a pyruvate decarboxylase), ii) the removal of the ldhA, frdA, and pflB genes that compete with the ethanol production route from pyruvate, and iii) the increase of ethanol tolerance by adaptive evolution.

### 4.2 Plasmid Assembly

Plasmids for ELP surface display were constructed from the base Addgene plasmid #186314. We replaced the displayed peptide with the fusion construct SpyTag-ELP-HisTag. Plasmids were cloned using Golden Gate Assembly^48^. Every plasmid was fully sequenced through Oxford Nanopore Technologies (service provided by Plasmidsaurus). Upon verification, plasmids were transformed into *E. coli* strains. Specific ELP sequences are reported in the Supplementary, Section 1.3.

### 4.3 Cell Growth

Engineered and wild-type *E. coli* strains were grown by inoculating single colonies into 3 mL of LB medium in 14 mL culture tubes. For the ethanologenic strain, LB medium was supplemented with α-lactose at a final concentration of 2 g/L. Cultures were grown overnight at 37 °C with shaking at 250 rpm. Overnight cultures were then diluted 1:100 into fresh medium and transferred to either 14 mL culture tubes (final volume: 7 mL) or 125 mL glass flasks (final volume: 50 mL). Cultures were grown to mid-log phase (OD_600_ = 0.4–0.6) and subsequently induced with either isopropyl β-d-1-thiogalactopyranoside (IPTG) or N-3-oxohexanoyl-L-homoserine lactone (HSL) at the indicated concentrations. Sedimentation assays, flow cytometry, and confocal microscopy were performed after 2 h of post-induction growth under shaking conditions. Imaging of macroscopic assemblies in flasks was conducted following overnight induction.

### 4.4 Confocal Microscopy

To verify the extracellular localization of ELPs, engineered cells were collected by centrifugation at 4,000 rpm for 1 min, resuspended in 1 mL of 0.01 M phosphate-buffered saline (PBS), and washed twice by repeated centrifugation and resuspension. Washed cells were incubated with 80 μg of SpyCatcher-GFP for 60 min at 30 °C to allow formation of covalent SpyTag-SpyCatcher bonds^19^. Samples were then washed three times with 1 mL of 0.01 M PBS and mounted between a slab of PBS agarose (1.5% w/v) and a glass coverslip. Imaging was performed using an Olympus FLUOVIEW FV3000 confocal laser scanning microscope. Images were acquired using FLUOVIEW software and analyzed with Fiji software.

### 4.5 Flow Cytometry-based GFP Quantification

Cells were incubated with SpyCatcher-GFP as described in Section 4.4 and subsequently washed three times with 1 mL of 0.01 M phosphate-buffered saline (PBS). For each sample, 10,000 events were acquired using a BD FACSCelesta flow cytometer. Data acquisition was performed with BD FACSDiva software, and data analysis was conducted using FlowJo software.

### 4.6 Viability Assay

To assess the viability of engineered and wild-type cells, propidium iodide (PI) staining was performed. Cells induced for 2 h were collected by centrifugation, resuspended in 1 mL of 0.01 M phosphate-buffered saline (PBS), and washed twice by repeated centrifugation and resuspension. An aliquot of 200 μL of the washed culture was transferred to a new 1.5 mL microcentrifuge tube for staining. Samples were incubated with 10 μL of propidium iodide (Thermo Fisher Scientific) on a Corning LSE platform rocker for 15 min to allow PI uptake by membrane-compromised cells. Stained cells were imaged by confocal microscopy as described in Section 4.4.

### 4.7 Sedimentation Assay

Sedimentation assays were performed as described in Trunk et al.^49^ Specifically, cells were grown in tubes, as described in Section 4.3. After 2 h of induction, cell cultures were incubated statically at 37 °C. Every hour, the OD_600_ was measured by sampling 200 µL of culture from the top of the tube and diluting it into 800 µL of fresh LB. Sedimentation curves were plotted in GraphPad Prism. All curves were normalized to the average OD_600_ value at t=0.

### 4.8 FloEc-ELMs Apparent Area Quantification

Flask-bottom images were analyzed to quantify the apparent area occupied by FloEc-ELMs. Images were processed in two stages: (i) detection of the circular flask-bottom region of interest (ROI) and (ii) segmentation of FloEc-ELM assemblies within the ROI. To ensure robustness to illumination and camera-dependent effects, images were analyzed in a device-independent color space. The ROI was identified based on the characteristic yellow background of the flask bottom using chromatic information, followed by morphological filtering and geometric selection of the most circular region. FloEc-ELMs within the ROI were segmented using a hybrid approach combining chromatic deviation from the background and enhancement of thin, branched, filament-like structures in the intensity domain. To minimize edge artifacts, segmentation was restricted to an inner ROI excluding the rim. Filamentous assemblies were detected using multi-scale line-enhancement and adaptive thresholding, followed by morphological reconstruction to recover entire branched structures while suppressing background noise. Apparent surface area of the FloEc-ELMs was quantified by converting segmented filament area from pixel units to physical units using an image-specific pixel-to-area scaling factor. This factor was derived from the detected flask-bottom ROI by assuming a known real diameter (67 mm), allowing filament area to be reported in mm² while compensating for small variations in ROI detection across images.

### 4.9 Microbial Fermentation

Pre-cultures of the ethanologenic strain W105Fe were initiated from single colonies and grown in 5 mL of LB Lennox medium supplemented with 40 g/L lactose at 37 °C and 220 rpm for 16 h. Lactose fermentations in rich medium were initiated by a 100-fold dilution of the preculture into 5 mL of LB Lennox containing 40 g/L lactose and 100 mM phosphate buffer (pH 7.0) in 50 mL conical tubes, followed by incubation at 37 °C and 220 rpm for 24 h. Where indicated, N-3-oxohexanoyl-L-homoserine lactone (HSL) was added 2 h after inoculation at a final concentration of 2 nM to induce FloEc-ELM assembly in the engineered W105Fe–ELP_10_-LYS strain.

Dairy waste fermentations were carried out using concentrated whey permeate (CWP) wastewater obtained from a whey valorization plant (Serum Italia S.p.A., Cazzago San Martino, Italy), containing more than 100 g/L lactose. The fermentation medium was prepared by diluting CWP twofold in deionized water and supplementing it with 150 mM piperazine-N,N′-bis(2-ethanesulfonic acid) (PIPES, pH 7.0) and 0.05% (w/v) ammonium sulphate as a nitrogen source. Fermentations were performed in a final volume of 5 mL in a 50-mL tube and inoculated using biomass recovered from 1.160-mL precultures.

Two biomass recovery strategies were evaluated before inoculation: (i) centrifugation at 4,000 rpm for 15 min, followed by resuspension of the bacterial pellet in fermentation medium; or (ii) sedimentation for 1.5 h at 37 °C, followed by decanting of the supernatant and use of the sedimented fraction (approximately 100 μL) for inoculation. Dairy waste fermentations were incubated at 37 °C and 220 rpm for 72 h. Growth conditions were based on previous works on ethanol production from dairy waste^43^.

In all fermentations, samples were collected at t = 0 and at the indicated timepoints (24 or 72 h), filtered through 0.2 μm membranes, and stored at −20 °C before quantification of fermentation products.

### 4.10 Quantification of Lactose and Ethanol Concentrations

A high-performance liquid chromatography system (LC-2000 HPLC, Jasco) was used to detect the residual lactose and produced ethanol at the end of fermentation experiments. An autosampler was used to inject 25 μL of samples, which were analyzed with a refractive index detector (RID, Shimadzu). The Supelco C-610H 30 cm × 7.8 mm column (59320-U, Sigma Aldrich), kept at 30°C, and the 0.1% H_3_PO_4_ mobile phase at the 0.5 mL/min flow rate were used for the analyses^47^.Quantifications were carried out from the measured chromatograms by the ChromNAV 2.0 software (Jasco). Ethanol titer (i.e., the concentration at the end of the fermentation) and conversion yield were used as fermentation performance indexes. The conversion yield was computed as a percentage of the maximum theoretical yield, considering that the maximum ethanol production corresponds to 4 molecules of ethanol per molecule of consumed lactose^47^.

### 4.11 Filtration Assay

Terminal fermentation cultures (100 mL) were transferred into 100-ml cylinders and left at 37°C for 1.5 h to allow for sedimentation. A 75-ml volume was then taken from the non-sedimented fraction and loaded into a 250-ml vacuum filtration unit with a 0.22-μm PES membrane (Biofil). Vacuum was applied by a diaphragm mini-pump (PM20405-86, VWR International GmbH) with a maximum pressure of 2.4 bar and a vacuum of 250 mbar. The time needed to filter the entire volume (75 ml) was recorded. Flow rate was computed as the volume divided by the overall filtration time.

### 4.12 Statistical Analysis

Comparisons between three or more groups were performed using the Brown-Forsythe and Welch ANOVA. ANOVAs had multiple comparisons post hoc analysis (Dunnett T3) performed. For comparisons among fewer than three groups, a two-tailed Welch’s t-test was used. In all cases, a p-value (p) cutoff of 0.05 was used to identify statistically significant differences. In the graphs, **** = p<0.0001, ** = p<0.01, and * = p<0.05.

## Supporting Information

The following file is available free of charge.

- Tables describing the strains and ELP plasmids cloned and used in the experiments; main sequences utilized in the paper; Western blot analysis of two surface-displayed ELP constructs; fluorescent flow cytometry for quantification of ELP constructs in the different strains used in the studies; confocal images of propidium iodide stained cells to understand cell health across WT and ELP containing cells; flask photographs to discern macroscopic material formed from the various ELPs; figure describing the sedimentation of the W105Fe and W105Fe ELP10-LYS cells after dilutions at different initial cell densities; photographs of W105Fe and W105Fe ELP10-LYS cells after collection through standard centrifugation or sedimentation methods (DOCX)

## Author Contributions

The manuscript was written through the contributions of all authors. All authors have given approval to the final version of the manuscript.

## Funding Sources

Army Research Lab, W911NF-23-2-0040

NIH NIGMS, 1R35GM157144

MUR–M4C2 1.5 of PNRR, ECS00000036

## Acknowledgements

We thank Dr. Birthe Kjellerup (UMD) for her thoughtful feedback on this work and Michela Casanova (Univ. Pavia) for fermentation support. SM acknowledges the Army Research Lab (grant no. W911NF-23-2-0040) and the NIH NIGMS (grant no. 1R35GM157144) for supporting this study. These two funding sources supported part of the development and characterization of ELMs described in the first two result sections. PM and LP acknowledge MUR—M4C2 1.5 of PNRR (grant ECS00000036, NODES project).

## Abbreviations

ALA: Alanine
ELM: Engineered Living Material
ELP: Elastin Like Polypeptides
FloEc-ELMs: Flocculating *E. coli*-based Engineered Living Materials
GLU: Glutamic Acid
His-Tag: Histidine Tag
HSL: N-3-oxohexanoyl-L-homoserine lactone
IPTG: Isopropyl β-d-1-thiogalactopyranoside
LYS: Lysine
PI: Propidium Iodide
SC: SpyCatcher
ST: SpyTag.

