## Supplementary Information for "De novo Engineered Living Materials via Elastin-Like Polypeptide-Mediated Self-Assembly"

**1. Supplementary Methods**

***1.1 Strains used in this study***

The *E. coli* strains used in this study are outlined in the table below.

| **Strain Description** | **Strain Designation** | **Source** | **Genotype** |
| --- | --- | --- | --- |
| wild type | BL21 | NEB | *fhuA2 [lon] ompT gal (λ DE3) [dcm] ∆hsdS*  *λDE3= λsBamHIo ∆EcoRI-B int::(lacI::PlacUV5::T7 gene1) i21 ∆nin5* |
| ethanologenic | W105Fe | L. Pasotti *et al., 2023* | *E. coli W (DSM 1116) ΔldhA ΔpflB-focA frdAB::P_J105_-adhB-pdc*  ***Including adaptive evolution for higher ethanol tolerance |

***1.2 Plasmids used in this study***

| **Plasmid Name** | **Main Features** | **Use** | **Source** |
| --- | --- | --- | --- |
| pSB035 | pT5_PelB_Ala10_ST_EhaA_AmpR | ELP_10_-ALA display | This work |
| pSB066 | pT5_PelB_Glu10_ST_EhaA_AmpR | ELP_10_-GLU display | This work |
| pSB045 | pT5_PelB_Lys10_ST_EhaA_AmpR | ELP_10_-LYS display | This work |
| pDP005 | pLux_PelB_Ala10_ST_EhaA_LuxR_  pConIq_KanR | ELP_10_-Lys display with LuxR/  HSL inducible system suitable for biofermentation | This work |
| pSBCC001 | pCon_BBa_J23116_mScarlet_KanR | Constitutive red fluorescence - Confocal | This work |
| pSMCAF016 | pBAD_HisTag_TEV_SpyCatcher_AmpR | SpyCatcher-GFP production | S. Molinari *et al.*, 2022 |
| pSB023 | pBAD_HisTag_TEV_SpyTag_AmpR | SpyTag-GFP production | This work |

***1.3 ELP and Important Sequences used in this study***

*ELP_10_-ALA*

GTCCCAGGAGTGGGTGTACCTGGTGCAGGAGTGCCGGGTGTAGGCGTTCCTGGCGCTGGAGTCCCGGGGGCTGGAGTTCCCGGCGTGGGCGTTCCTGGTGCAGGCGTGCCGGGCGTCGGAGTGCCGGGAGCGGGTGTTCCGGGGGCCGGT

*ELP_10_-GLU*

GTCCCAGGAGTAGGCGTACCTGGTGTGGGAGTACCCGGTGAAGGTGTGCCGGGTGTGGGCGTACCGGGCGAAGGAGTTCCTGGTGTGGGCGTGCCTGGCGTTGGCGTTCCGGGCGAAGGTGTGCCAGGGGTTGGGGTGCCCGGTGAAGGT

*ELP_10_-LYS*

GTCCCAGGAGTGGGGGTCCCGGGGAAAGGGGTGCCCGGCGTTGGGGTCCCTGGTAAAGGCGTTCCTGGCAAAGGCGTGCCTGGCGTTGGGGTTCCTGGAAAAGGCGTACCCGGAGTAGGTGTTCCGGGTAAAGGTGTACCCGGGAAAGGT

*SpyTag003*

CGTGGTGTTCCGCACATTGTTATGGTTGATGCGTATAAACGTTATAAA

*SpyCatcher003*

ATGGTTGATACCTTATCAGGTTTATCAAGTGAGCAAGGTCAGTCCGGTGATATGACAATTGAAGAAGATAGTGCTACCCATATTAAATTCTCAAAACGTGATGAGGACGGCAAAGAGTTAGCTGGTGCAACTATGGAGTTGCGTGATTCATCTGGTAAAACTATTAGTACATGGATTTCAGATGGACAAGTGAAAGATTTCTACCTGTATCCAGGAAAATATACATTTGTCGAAACCGCAGCACCAGACGGTTATGAGGTAGCAACTGCTATTACCTTTACAGTTAATGAGCAAGGTCAGGTTACTGTAAATGGCAAAGCAACTAAAGGTGACGCTCATATT

*GFP*

ATGCGTAAAGGCGAAGAGCTGTTCACTGGTGTCGTCCCTATTCTGGTGGAACTGGATGGTGATGTCAACGGTCATAAGTTTTCCGTGCGTGGCGAGGGTGAAGGTGACGCAACTAATGGTAAACTGACGCTGAAGTTCATCTGTACTACTGGTAAACTGCCGGTTCCTTGGCCGACTCTGGTAACGACGCTGACTTATGGTGTTCAGTGCTTTGCTCGTTATCCGGACCATATGAAGCAGCATGACTTCTTCAAGTCCGCCATGCCGGAAGGCTATGTGCAGGAACGCACGATTTCCTTTAAGGATGACGGCACGTACAAAACGCGTGCGGAAGTGAAATTTGAAGGCGATACCCTGGTAAACCGCATTGAGCTGAAAGGCATTGACTTTAAAGAAGACGGCAATATCCTGGGCCATAAGCTGGAATACAATTTTAACAGCCACAATGTTTACATCACCGCCGATAAACAAAAAAATGGCATTAAAGCGAATTTTAAAATTCGCCACAACGTGGAGGATGGCAGCGTGCAGCTGGCTGATCACTACCAGCAAAACACTCCAATCGGTGATGGTCCTGTTCTGCTGCCAGACAATCACTATCTGAGCACGCAAAGCGTTCTGTCTAAAGATCCGAACGAGAAACGCGATCATATGGTTCTGCTGGAGTTCGTAACCGCAGCGGGCATCACGCATGGTATGGATGAACTGTACAAATAA

*mScarlet*

GTGAGTAAAGGAGAAGCTGTGATTAAAGAGTTCATGCGCTTCAAAGTTCACATGGAGGGTTCTATGAACGGTCACGAGTTCGAGATCGAAGGCGAAGGCGAGGGCCGTCCGTATGAAGGCACCCAGACCGCCAAACTGAAAGTGACTAAAGGCGGCCCGCTGCCTTTTTCCTGGGACATCCTGAGCCCGCAATTTATGTACGGTTCTAGGGCGTTCATCAAACACCCAGCGGATATCCCGGACTATTATAAGCAGTCTTTTCCGGAAGGTTTCAAGTGGGAACGCGTAATGAATTTTGAAGATGGTGGTGCCGTGACCGTCACTCAGGACACCTCCCTGGAGGATGGCACCCTGATCTATAAAGTTAAACTGCGTGGTACTAATTTTCCACCTGATGGCCCGGTGATGCAGAAAAAGACGATGGGTTGGGAGGCGTCTACCGAACGCTTGTATCCGGAAGATGGTGTGCTGAAAGGCGACATTAAAATGGCCCTGCGCCTGAAAGATGGCGGTCGCTATCTGGCTGACTTCAAAACCACGTACAAAGCCAAGAAACCTGTGCAGATGCCTGGCGCGTACAATGTGGACCGCAAACTGGACATCACCTCTCATAATGAAGATTATACGGTGGTAGAGCAATATGAGCGCTCCGAGGGTCGTCATTCTACCGGTGGCATGGATGAACTATACAAA

***1.4 Primers used to assemble plasmids***

*Primers used to assemble pSB035_Ala10*

SB220 - CAGGTCTCCCGGTCGTGGTGTTCCGCACATT

SB221 - CAGGTCTCCGGACTCCTGAGTGATGGTGATGG

*Primers used to assemble pSB066_GlutAcid10*

SB402 – AAGGTCTCCTGGTGAAGGAGTACCCGGTGAAGG

SB403 – TGGGTCTCCCCTGGCACACCTTCGCCCGG

SB400 – AAGGTCTCCCAGGGGAAGGGGTGCCCGGT

SB401 - AAGGTCTCCACCAGGTACGCCTACTCCTGGGA

*Primers used to assemble pSB045_Lys10*

SB252 - CAGGTCTCCGTCGTGGTGTTCCGCACATT

SB253 - CAGGTCTCCACTCCTGAGTGATGGTGATGG

*Primers used to assemble pDP005_Plux_Lys10*

DP027 - GAGGTCTCGAATGGCGACGAACAATAAGG

DP028 - GAGGTCTCGTTAGAAACGATCCTCCGCA

DP029 - GAGGTCTCGCTAATGAAATACCTGCTGCCG

DP030 - GAGGTCTCGCATTAGAATTGCCACTTAATGC

*Primers used to assemble pSBCC001*

CC001-CTGGTCTCCAATGGCGACGAACAATAAGG

CC002-CTGGTCTCCATCATTAGAAACGATCCTCCGC

CC003-CTGGTCTCGTGATGGTGAGTAAAGGAGAAGC

CC004-CTGGTCTCGCATTATCATTTGTATAGTTCATCCATGC

*Primers used to assemble pSB023_SpyTag003_GFP*

SB090-GTGGTCTCCAAAGGTGGAGGTTCAGGCGGT

SB091-GTGGTCTCCCACGGCCCTGAAAATACAGGTTTTCG

SB092-GTGGTCTCGCGTGGTGTTCCGCACATT

SB093-GTGGTCTCGCTTTATAACGTTTATACGCATCAACC

**2. Supplementary Figures**

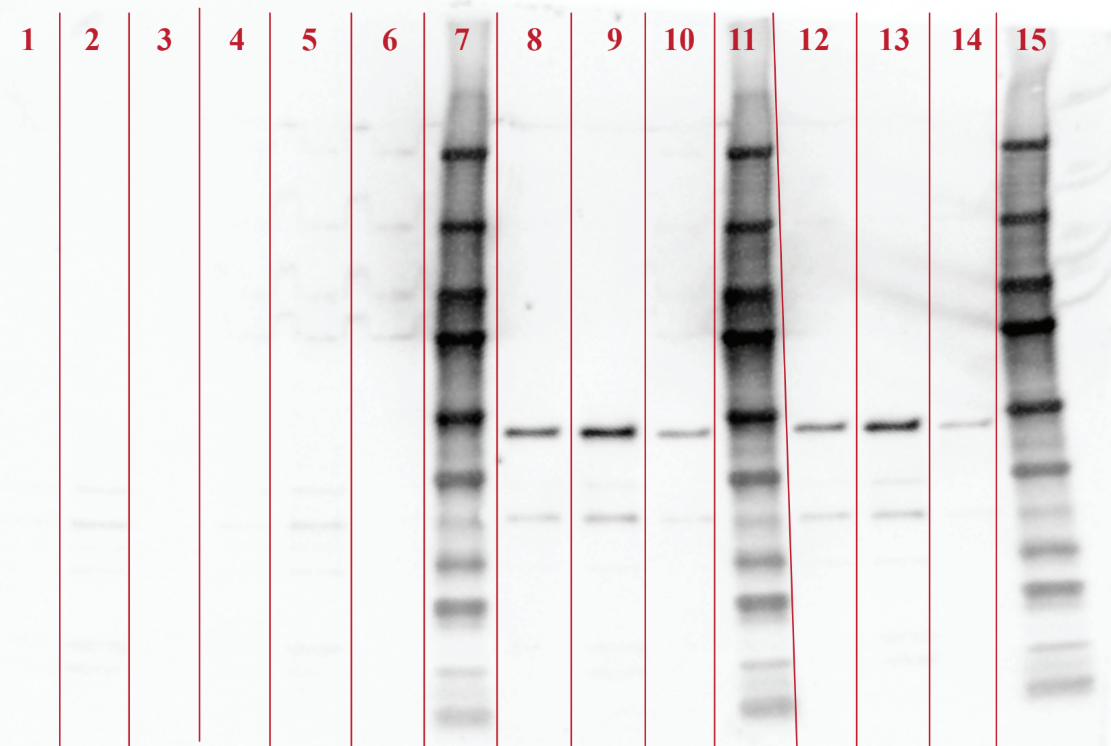

**Supplementary Figure 1. Western blot analysis of surface-displayed ELP constructs.** Western blot showing detection of ELP_10_-ALA induced with 0 µM IPTG (lanes 1-3) and 25 µM IPTG (lanes 8-10), and ELP_10_-LYS induced at 0 µM IPTG (lanes 4-6) and 25 µM IPTG (lanes 12-14). For each condition, lanes correspond (from left to right) to whole-cell lysate, membrane fraction, and cytosolic fraction. Upon induction, both constructs show clear enrichment in the membrane fraction (lanes 9 and 13), consistent with successful surface display. The expected molecular weight of the displayed ELP constructs is 46.17 kDa, and all detected bands migrate at the anticipated position just below the 50 kDa marker (Precision Plus Western C). Detection was performed following overnight induction using anti-HisTag antibodies (Thermo Fisher Scientific, PA1-983B).

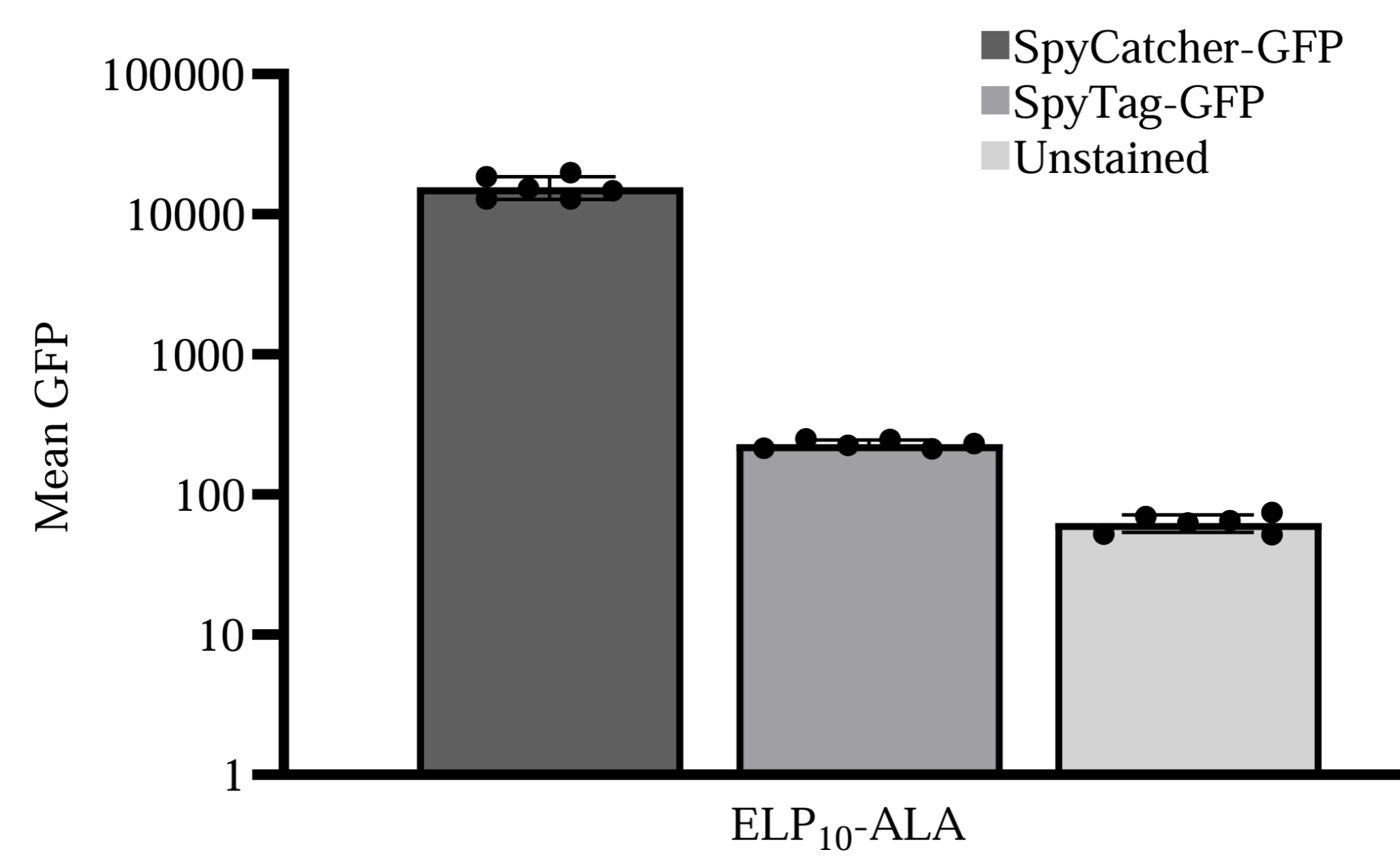

**Supplementary Figure 2. Quantitative flow cytometry of SpyCatcher-GFP-stained cells displaying ELP_10_-ALA.** (Left) Mean GFP detection for ELP_10_–ALA stained with SpyCatcher-GFP after 2 hours of induction with 25 µM IPTG. Error bars are centered on the mean value and represent 95% confidence intervals of six samples.

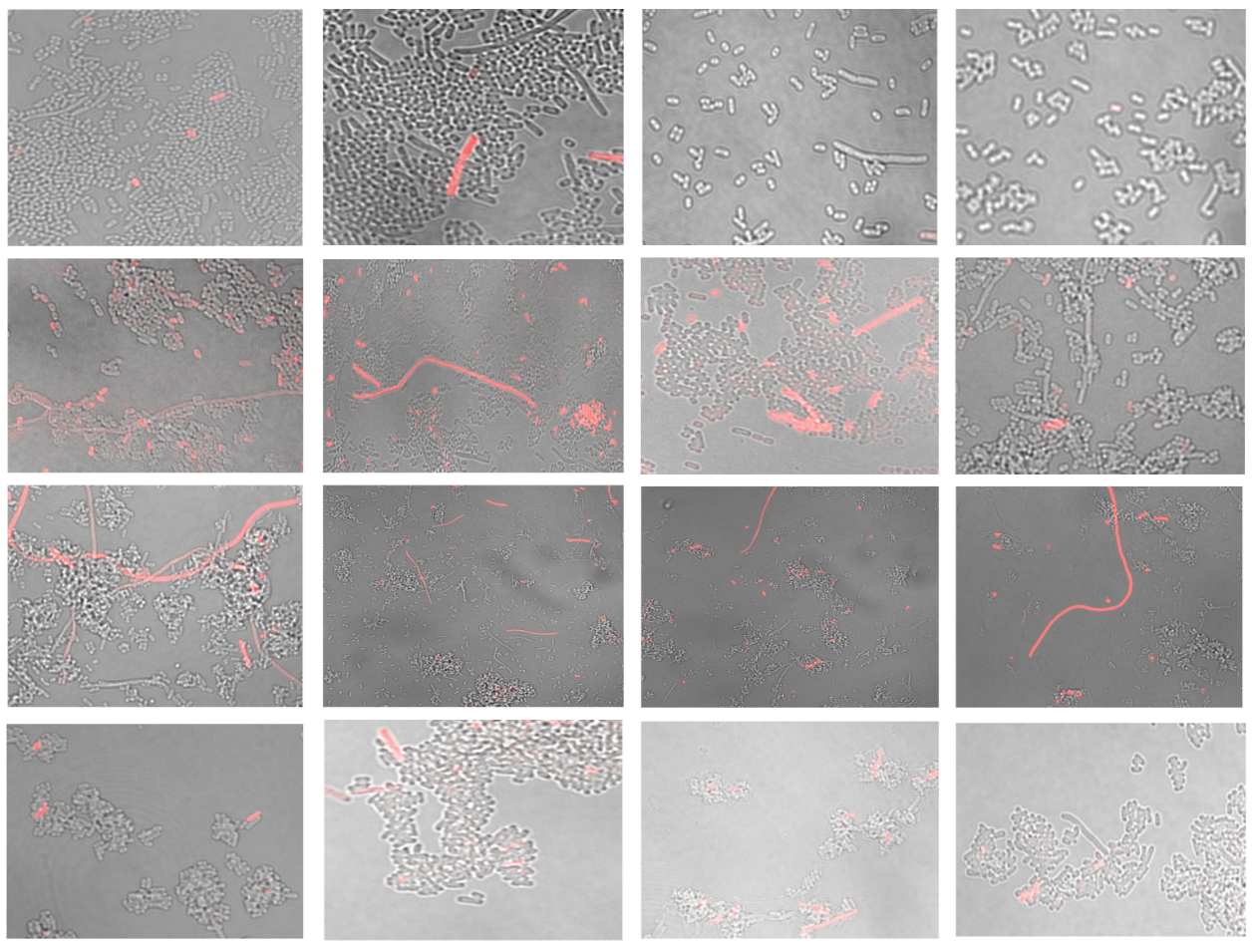

BL21 WT

ELP_10__ALA

**Supplementary Figure 3. Confocal microscopy of PI-stained cells.** Representative images of confocal microscopy for BL21 WT cells and ELP_10_–ALA displaying cells after being induced with IPTG as described in the Methods section of the main text and stained with PI.

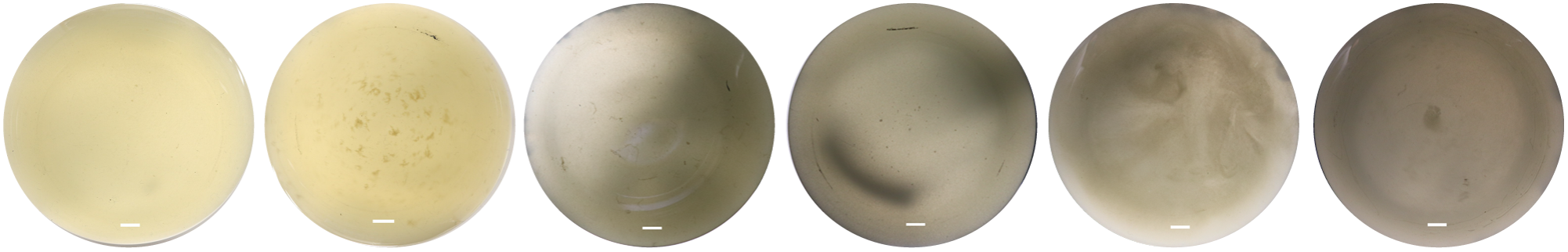

**Supplementary Figure 4. Flask panel from overnight induction of ELP_10_-ALA.** Photographs of the inconsistent FloEc-ELMs produced from ELP_10_-ALA. Photographs taken from the bottom of culture flasks of cell lines grown overnight under shaking conditions and induced with 25µM IPTG. The scale bars represent 0.5 cm.

**Supplementary Figure 5.** **Full fluorescent quantitative flow cytometry of displayed ELP_10_-ALA_,_ ELP_10_-GLU_,_ and ELP_10_-LYS.** Mean GFP detection for each displayed ELP constructs after 2 hours of induction. In addition to SpyCatcher-GFP staining, SpyTag-GFP and unstained groups are reported as negative control (the SpyTag-GFP values were used for normalization in Fig. 2A). Error bars are centered on the mean value and represent 95% confidence intervals of six samples.
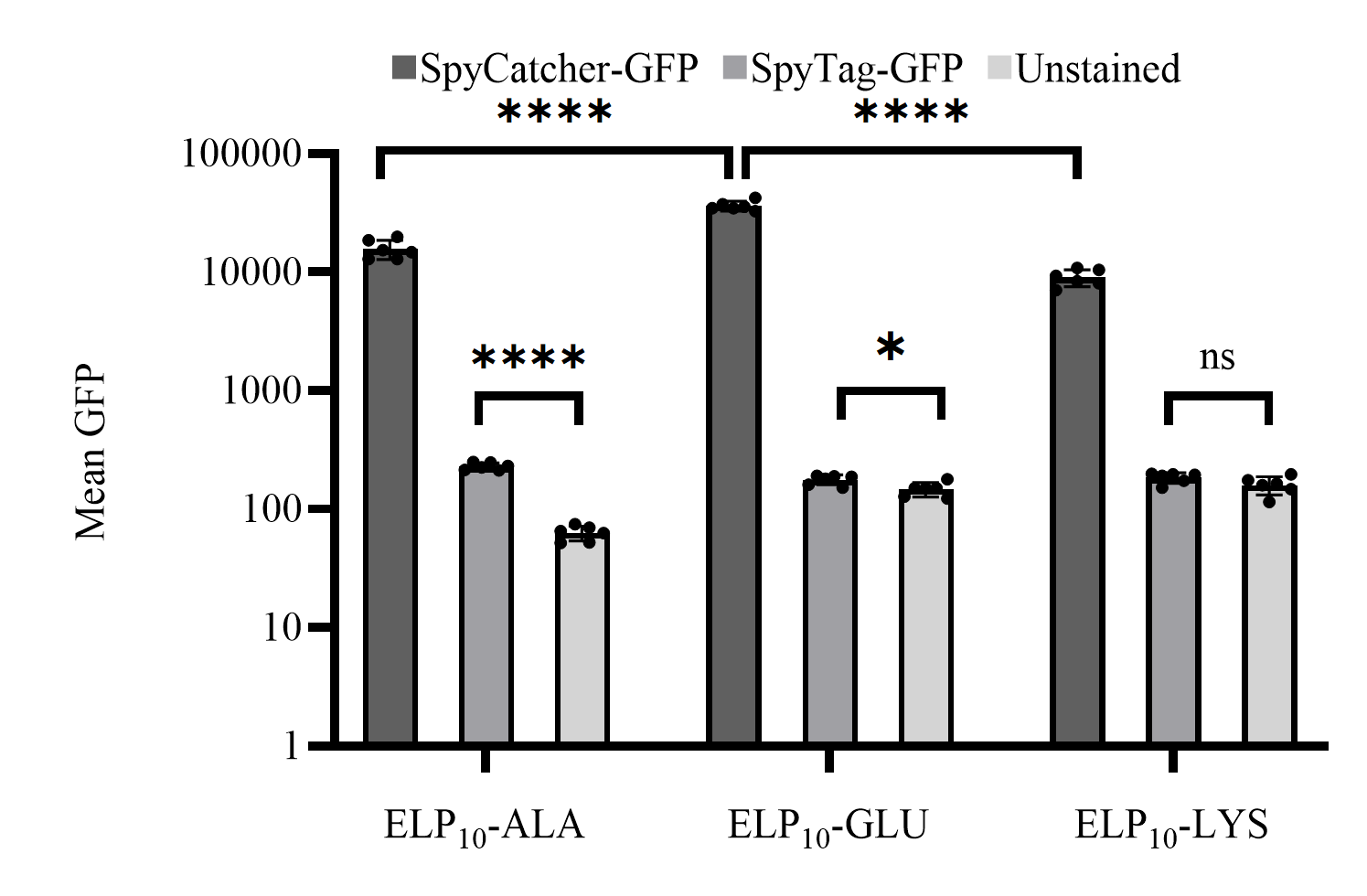

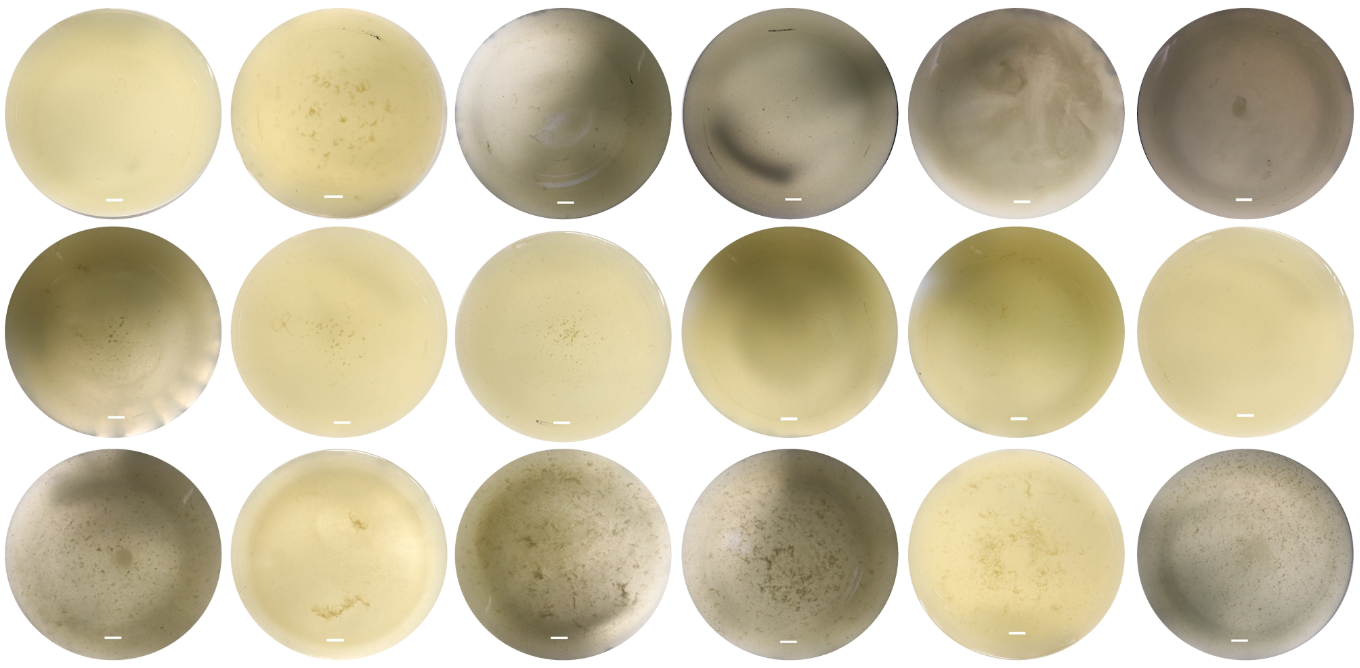

**Supplementary Figure 6. Photographs of FloEc-ELMs**. Photographs taken from the bottom of culture flasks of cell lines grown overnight under shaking conditions and induced with 25µM IPTG – Top: ELP_10_-ALA (same as Supplementary Fig. 4, copied here for direct comparison); Middle: ELP_10_-GLU. Bottom: ELP_10_-LYS. The scale bars represent 0.5 cm.

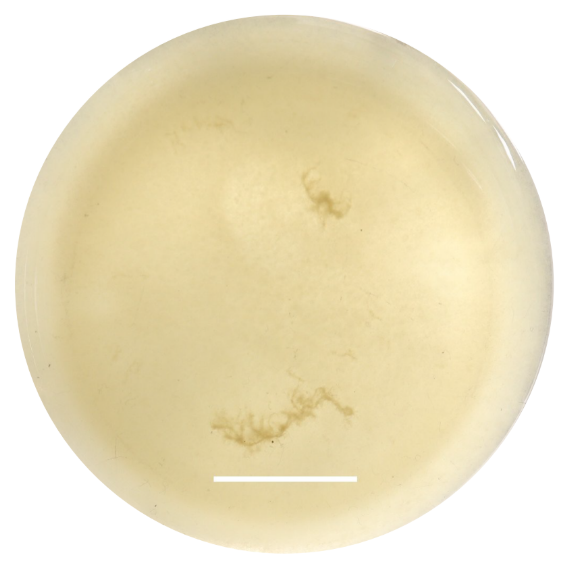

**Supplementary Figure 7. Flask image portraying the largest FloEc-ELM produced from cells displaying ELP_10_-LYS.** The depicted FloEc-ELM was formed overnight from cells induced with 25µM IPTG. The pictured bar is 1.75 cm long.

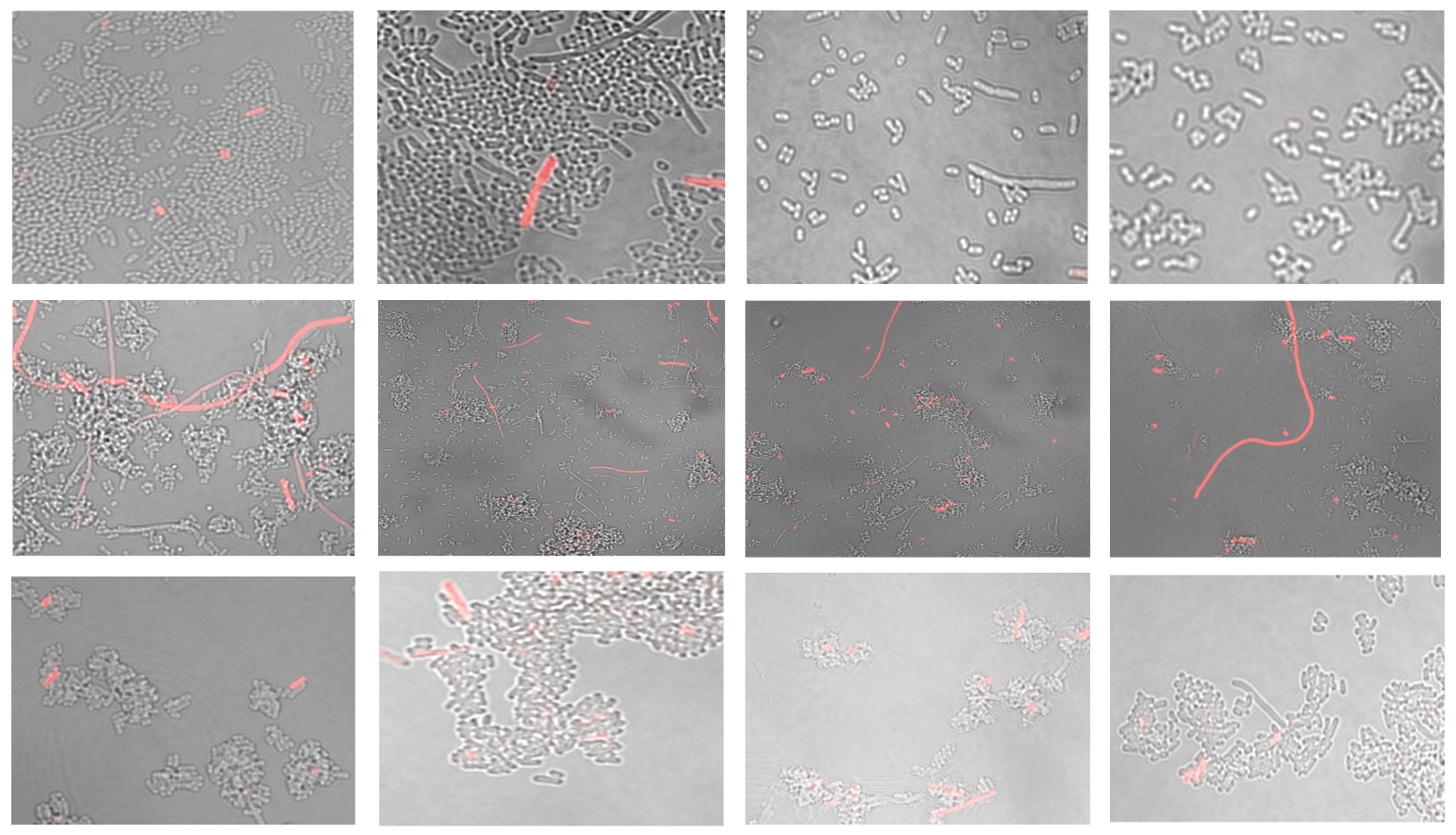

ELP_10_-GLU

ELP_10_-LYS

**Supplementary Figure 8. Confocal microscopy of PI-stained ELP_10_-GLU and ELP_10_-LYS cells.** Representative confocal microscopy images of BL21 wild-type cells and ELP_10_-ALA-engineered cells following IPTG induction, as described in the Methods, and stained with propidium iodide (PI).

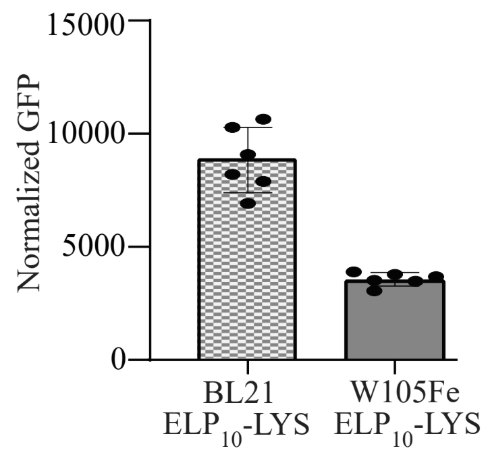

**Supplementary Figure 9. Flow cytometry of the ELP_10_-LYS-displaying cells in both BL21 and W105Fe strains stained with SpyCatcher-GFP.** At 2 nM HSL induction, W105Fe–ELP_10_-LYS cells displayed lower surface fluorescence than their BL21 counterparts induced with 25 µM IPTG, albeit within the same order of magnitude. Error bars are centered on the mean value and represent 95% confidence intervals of six samples.

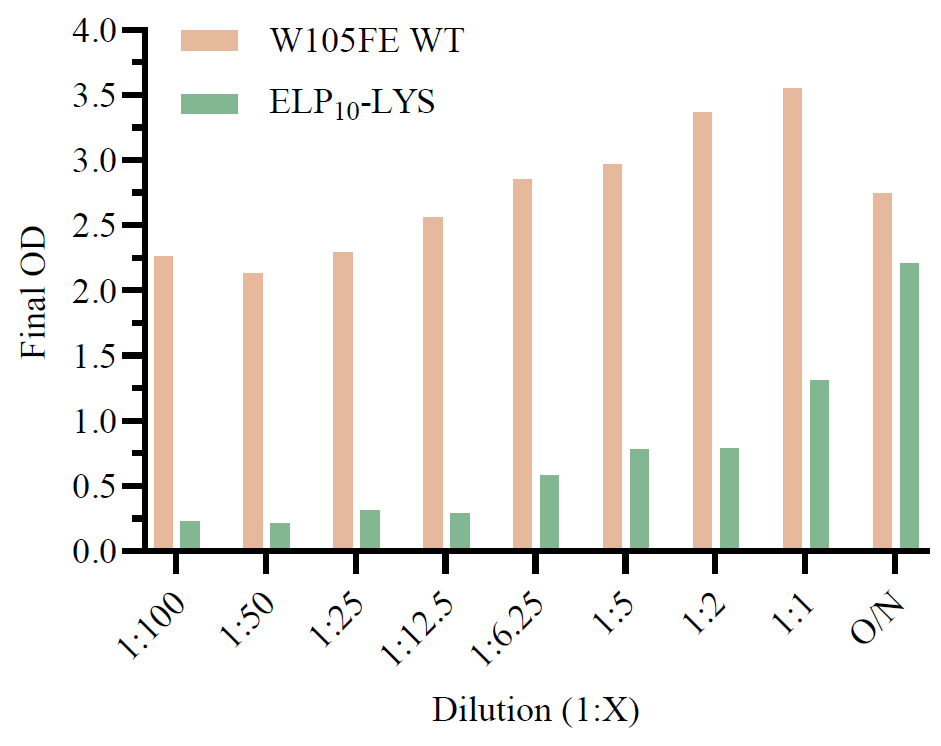

**Supplementary Figure 10. Sedimentation assay of W105Fe base and W105Fe ELP_10_-LYS strain diluted at different initial cell densities.** OD_600_ values after 4 h of static incubation of the W105Fe WT and W105Fe-ELP_10_-LYS strain. Cells from an overnight (O/N) preculture were diluted as indicated in the X-axis, in fresh LB + 2 g/L lactose. Cultures were induced with 2 nM HSL after 1.7 h from dilution (time needed to reach the exponential phase after a 100-fold dilution). At the time of HSL addition, OD_600_ values were comparable between the two strains for each dilution level (data not shown).

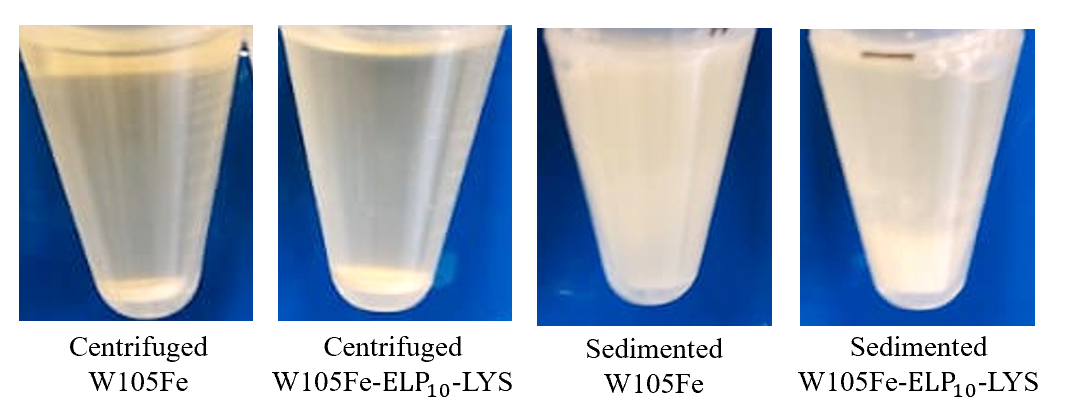

**Supplementary Figure 11. Images of the W105Fe and W105Fe-ELP_10_-LYS following centrifugation and sedimentation methods of collecting pre-culture.** Notably, the W105Fe-ELP_10_-LYS strain exhibited prominent deposition (Sedimented W105Fe-ELP_10_-LYS) not visible in the base strain (Sedimented W105Fe).
